# COVID19-vaccination affects breath methane dynamics

**DOI:** 10.1101/2022.07.27.501717

**Authors:** Daniela Polag, Frank Keppler

## Abstract

Methane (CH_4_) is well known as a component in the exhaled breath of humans. It has been assumed for a long time that formation of CH_4_ in humans occurs exclusively by anaerobic microbial activity (methanogenesis) in the gastrointestinal tract. A fraction of the produced CH_4_ is excreted via the lungs and can then be detected in the breath. However, recent studies challenge this view by showing that CH_4_ might also be produced endogenously in cells by oxidative-reductive stress reactions. Thus, an increased and fluctuating level of breath CH_4_ compared to the base level of an individual might also indicate enhanced oxidative stress levels. Thus, monitoring breath CH_4_ levels might have great potential for ‘in vivo’ diagnostics.

Generally, vaccines generate a strong immune response including the production of pro-inflammatory cytokines. To evaluate the effect from current vaccines against COVID-19 on breath CH_4_ dynamics, breath CH_4_ was monitored from 12 subjects prior and after the injection of several COVID-vaccines. Prior to COVID-19 vaccination the concentration of breath CH_4_ was frequently measured by gas chromatograph flame ionization detection (GC-FID, with analytical precision better than 10 parts per billion, ppbv) to obtain the individual variation range of breath CH_4_ for each subject. Following vaccination, CH_4_ breath samples were collected at high frequency for a period of 14 days.

All subjects monitored showed a strong response in breath CH_4_ release within 1 to 72 hours after vaccination including shifts and high fluctuations with maximum peaks showing a factor of up to ±100 compared to base values. Thus, it is highly likely that the observed changes in breath CH_4_ are coupled to immune responses following Covid-19 vaccination. These preliminary results strongly support the hypothesis that non-microbial methane liberation and utilisation in the human body might be also linked to cellular processes and stress responses independent of classical microbial methanogenesis. Thus, CH_4_ might be used as a breath biomarker for specific immune responses and individual immune states.

## 1 Introduction

Since the last decade, there has been a rising interest in clinical volatile biomarkers for specific pathogen diagnostics, characterisation of physical and cellular state, and monitoring with respect to baseline variations (Aronson & Ferner, 2017). Breath testing as a rapid, cheap, point-of-care, and non-invasive diagnostic tool is known since some time (Manolis, 1983). Nowadays, specific volatile organic compounds (VOCs) or VOC profiles in excreted breath are considered as promising candidate biomarkers used to detect various functional diseases and pathogenic infections (Das & Pal, 2020). As human breath contains more than 3500 VOCs of often unknown sources breath research it is not only an analytical challenge but limited knowledge exists about the significance and underlying mechanisms of single VOC compounds and VOC profiles regarding pathogen diagnostics (Phillips, et al., 2010), (Hong-Geller & Adikari, 2018). Breath biomarkers simple in measurement and interpretation as well as unambiguously identifying deviations from baseline are the subject of current research (Nakhleh, et al., 2017), (Gaude, et al., 2019), (Ghosh, et al., 2021).

In recent years, CH_4_ as one of the simplest carbon-based VOC-molecules and a frequent component in the exhaled breath of humans has been considered as a potential biomarker providing information on microbial activity in the gastrointestinal tract (Di Stefano & Corazza, 2009), (De Lacy Costello, et al., 2013), (Bang, et al., 2014), (Ghavami, et al., 2018). Methanogens such as *Methanobrevibacter smithii* and *Methanosphaera stadtmanae* are usually present in the distal part of the colon where they metabolize hydrogen (H_2_) and methanol to form CH_4_ (Flourie, et al., 1991), (Mihajlovski, et al., 2010). A fraction of the CH_4_ produced is excreted via the lungs and can be detected by specific gas analysis (Mürtz & Hering, 2008), (Dryahina, et al., 2010), (Tuboly, et al., 2013).

The proportion of CH_4_ in exhaled breath has been investigated in numerous studies. However, most of these studies only differentiate between breath CH_4_ producers and breath CH_4_ non-producers (McKay, et al., 1985), (Bolin, et al., 1996), (Kinoyama, et al., 2006). Subjects are defined as methane producers when they contain at least 1 part per million by volume (ppmv) above background level in their breath samples. Based on this definition, approximately 25% of people can be identified as CH_4_ producers (Bond, et al., 1971), (Peled, et al., 1987). However, recent high-precision analysis results suggested a modification of the definition range of breath CH_4_ producers (Keppler, et al., 2016). Based on their study they suggested to have three categories: i) low emitters/light absorbers showing a range of methane values slightly above (< 1 ppmv) and below background values, ii) medium emitters with values between 1-4 ppmv above background and iii) high emitters with values above 4 ppmv.

Generally, CH_4_ producing status has been the subject of numerous studies evaluating relationships with age (Hopkins, et al., 2002), (Polag, et al., 2014), ethnic background (Pitt, et al., 1980), (Santos-Mello, et al., 2012), gender (Triantafyllou, et al., 2014), (Polag, et al., 2014), exercise status (Szabó, et al., 2015) and various gastrointestinal diseases (Montes, et al., 1993), (Conway de Macario & Macario, 2009), (Hwang, et al., 2010), (Roccarina, et al., 2010), (Kunkel, et al., 2011), (Furnari, et al., 2012). Moreover, it was found that with increasing age the percentage of high breath CH_4_ emitters continuously increases (Polag, et al., 2014) but also that person-specific breath CH_4_ vary drastically over short times, i.e. high emitters change to low emitters and vice versa (Polag & Keppler, 2018). Results from long term monitoring of breath CH_4_ showed that abrupt deviations in breath CH_4_ levels from baseline might be related to changes in the physiological state of a person including immune reactions and inflammatory processes. These findings support the hypothesis that next to CH_4_ production by obligate anaerobic methanogens a portion of CH_4_ might also be produced under aerobic conditions on a cellular level without the contribution of microorganisms (Tuboly, et al., 2013), (Boros, et al., 2015), (Tuboly, et al., 2017). In this context, it was suggested that CH_4_ has a bioactive role in cellular physiology of eukaryotes and might be applied as a marker of oxido-reductive stress (Boros & Keppler, 2018), (Boros & Keppler, 2019).

Alternative pathways to microbially induced methanogenesis under aerobic conditions in eukaryotic cells were first described by Keppler et al. (Keppler, et al., 2006), (Keppler, et al., 2009). They observed that terrestrial plants were able to produce CH_4_ under oxic conditions, and excluded the contribution by obligate anaerobic methanogens. Further studies with fungi, mosses, and algae (Lenhart, et al., 2012), (Lenhart, et al., 2015), (Lenhart, et al., 2016) unambiguously demonstrated that CH_4_ production in eukaryotes is not restricted to endogenous microbial processes. A recent publication by (Ernst, et al., 2022) who studied CH_4_ formation in over 30 model organisms, ranging from eukaryotes (plant cells and several human cell lines), archaea and bacteria proposed a common mechanism based on reactive oxygen species (ROS) generated by oxidative stress and the interaction with iron and methyl donors. Following this discovery, it has been proposed that monitoring CH_4_ as an indicator for ROS-driven processes might be a promising approach with associated diagnostic potential (Keppler et al., 2022). The human immune system is of particular interest regarding immune responses based on infection and vaccination influenced by a large number of individual factors (Zimmermann & Curtis, 2019). Various parameters are used to estimate immune response using high-throughput ‘omic’ approaches including genes, mRNA, proteins, and metabolites (Pulendran & Davis, 2020) but effort and complexity are high. Thus, the need for a parameter that is easy to measure and interpret is desirable.

Vaccination represent an induced perturbation of the immune system with the possibility to monitor immune responses as deviations from baseline biomarkers (Pulendran & Davis, 2020). Influenza vaccination for example is known to have an effect on oxidative stress products in breath (Hayney & Buck, 2002), (Phillips, et al., 2010).

In this study, we evaluated the effect from current vaccines against COVID-19 on breath CH_4_ dynamics to further validate the occurrence of CH_4_ as a biomarker for oxidative stress and its potential indication of individual immune states. We took advantage of the current high rates of COVID-19-vaccinations. Thus, we carried out an in-house study where CH_4_ breath levels were monitored for 12 subjects prior and after the first, second and third injection of several COVID-vaccines.

## 2 Material and methods

The effect from COVID-19 vaccination on breath CH_4_ mixing ratios was studied for 12 subjects, 6 males and 6 females between 23 and 61 years obtaining different COVID-19 vaccines. All subjects were volunteers from the Institute of Earth Sciences at Heidelberg University (Germany). Body mass indices (BMI’s) of the subjects were in a normal range (18.5 to 25) and no prescribed medication or drug intake was reported prior to commencement of the experiment.

For breath sample collections, 1l Tedlar bags were used. The procedure for breath CH_4_ sampling was always carried out in a similar manner. During sample collection the subjects breathed normally, stopped breathing for 5sec and then completely filled the Tedlar-bag with expired air.

For CH_4_ measurement a sample of 6 ml was removed from the Tedlar-bag using a gas tight syringe and analyzed using a gas chromatograph (column: 2m, ⌀ =3.175mm (i.d.) high-grade steel tube packed with molecular sieve 5A 60/80 mesh from Supelco) equipped with a flame ionization detector (FID) (Shimadzu GC-14B). Before entering the analytical system, the breath gas was passed through a chemical trap filled with Dryerite® (anhydrous CaSO_4_) to remove water. Quantification of CH_4_ was performed by direct comparison of peak area with that of two reference standards containing 8.905 and 1.835 ppmv CH_4_ air. Analytical precision was better than 10 ppbv and individual CH_4_ mixing ratios usually showed a 10% variability during a regular day.

For every breath sample an additional control sample (background air) was collected to determine the fraction of the endogenous CH_4_ mixing ratio. Therefore, the background air or inhaled air was subtracted from the measured breath sample to obtain endogenous CH_4_ production/consumption.

Each subject was his or her own control since breath CH_4_ was monitored daily to weekly prior to vaccination to resolve baseline variations which showed an overall range between -102 ppbv to 65000 ppbv below/above background. Following vaccination, breath CH_4_ samples were collected in high-resolution (hours to days) showing an overall breath CH_4_ range which varied between -1000 ppbv to 77000 ppbv.

Table. 1 provides an overview of the key data of the study showing information on subjects’ CH_4_ producing status, gender, age at the time of first COVID-19 vaccination, and administered COVID19-vaccines including physical vaccine response and p-values according to the vaccine effect.

**Tab.1:** Information on subjects’ CH4 producing status, gender, age, and administered COVID19-vaccines including vaccine response (+: physical discomfort, -: none) and p-value of vaccine effect (*: F-test, **: t-test, NA: data set too low for testing).

| ID | CH<sub>4</sub> status | sex | age | vaccine 1 | p-value v1 | vaccine 2 | p-value v2 | vaccine 3 | P-value v3 |
| --- | --- | --- | --- | --- | --- | --- | --- | --- | --- |
| 1 | low | F | 25 | BioNtech (-) | 0.0006* | BioNtech (-) | NA | BioNtech (-) | 0.045* |
| 4 | low | M | 52 | Astra (+) | 1·10<sup>-11</sup>* | BioNtech (-) | 1·10<sup>-7</sup>* | Moderna (+) | 0.018* |
| 8 | low | M | 30 | BioNtech (-) | 0.0004* | BioNtech (-) | 0.299*; 0.538** | - | - |
| 11 | low | M | 28 | Johnson (++) | 0.273*; 0.139** | Moderna (-) | NA | - | - |
| 3 | low | F | 32 | BioNtech (-) | 0.97*; 0.395** | BioNtech (-) | NA | BioNtech (-) | 0.2*; 0.5** |
| 9 | low | F | 23 | BioNtech (-) | 0.0176* | BioNtech (+) | NA | - | - |
| 10 | low/medium | M | 32 | BioNtech (-) | 1·10<sup>-6</sup>* | BioNtech (+) | 0.387*; 0.81** | - | - |
| 6 | medium | M | 54 | BioNtech (+) | 0.089*; 3·10<sup>-9</sup>** | BioNtech (+) | NA | Moderna (+) | 0.975*; 3·10<sup>-5</sup>** |
| 2 | medium | M | 61 | Astra (-) | 4·10<sup>-6</sup>* | Astra (-) | 2·10<sup>-6</sup>* | Moderna (-) | 0.5*; 0.0011** |
| 5 | high | F | 44 | Moderna (-) | 9·10<sup>-5</sup>* | Moderna (+) | NA | Moderna (+) | 0.17*; 0.066** |
| 7 | high | F | 28 | BioNtech (-) | 2·10<sup>-11</sup>* | BioNtech (++) | NA | BioNtech (++) | NA |
| 12 | high | M | 53 | Moderna (-) | 0.197*; 0.128** | Moderna (+) | NA | Moderna (+) | 0.388*; 0.314** |

As established by Keppler et al. (Keppler, et al., 2016), CH_4_ producing status was classified as low (<1ppmv above background), medium (1-4ppmv above background) and high (>4ppmv above background). In total, six subjects were classified as low emitters, three as medium emitters and three as high emitters.

To validate the significance of vaccine reaction, in a first step the variances of CH_4_ values were compared before and after vaccination by a two-sided paired F-test with α = 5%. A minimum of 8 time points before and after vaccination were used as test condition. In case of confirmation of the zero-hypothesis (p > 0.05), in a second step the mean values of CH_4_ values were compared before and after vaccination by a two-sided paired t-test with α = 5%. The p-values of the associated tests are included in Table 1.

## 3 Results

All of the investigated subjects (Tab.1) showed a drastically effect on breath CH_4_ dynamics (Fig. 1) after COVID-19 vaccination. Compared to the baseline values prior to vaccination, a shift and/or a high variability of breath CH_4_ values was observed within the first few hours to days following vaccination. Figure 1 shows the individual endogenous breath CH_4_ production within 0 to 14 days following the first, second and, if available, third vaccine administration. As breath CH_4_ values were corrected for background CH_4_ composition, positive CH_4_ values indicate endogenous CH_4_ production whereas negative values represent endogenous CH_4_ consumption. The 95% confidence intervals of base values prior to vaccinations are included in Figure 1.

**Fig. 1:**
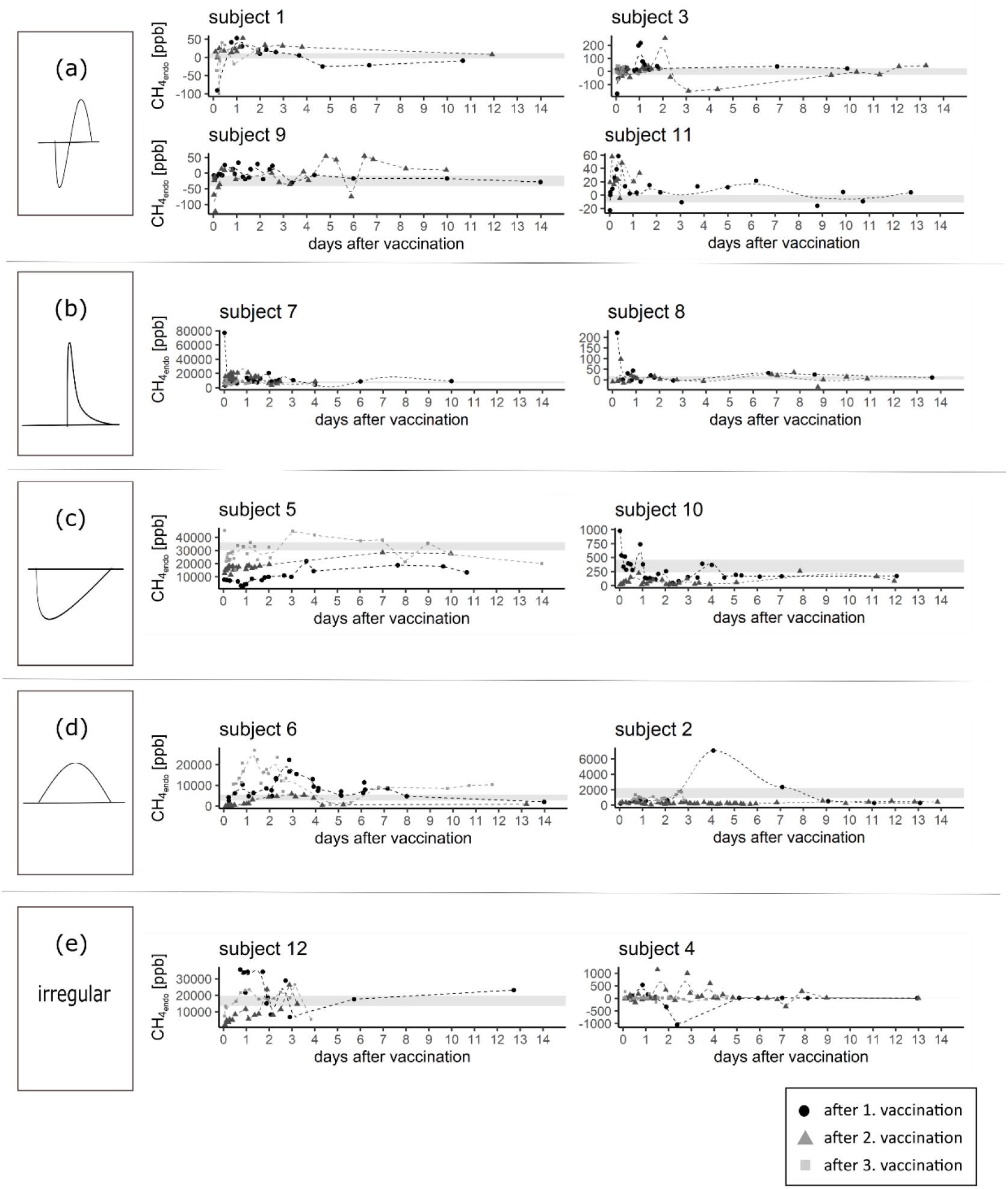
Endogenous breath CH_4 values_ (CH_4 endo_ = CH_4 breath_ – CH_4 background_) for the first 14 days following the first, second and, if available, the third COVID-19 vaccination (t=0) for the 12 subjects studied here. Grey shaded rectangles represent the 95% confidence interval of base CH_4 endo_ values monitored prior to vaccination and 4 weeks after vaccination, respectively. Breath CH4 dynamics are ordered by the resulting pattern: a) short-term dynamics with a negative initial peak, b) short-term dynamics with a positive initial peak, c) long-term dynamics with an initial decrease, d) long-term dynamics with an initial increase, e) irregular patterns. Uncertainty of CH4 values lie in a range of 10% (intraday fluctuation range). Please note the large differences of scales (y-axis) between the different subjects.

Here we distinguish between five CH_4_ breath patterns divided by (i) the direction of the initial shift (positive = patterns (b), (d) or negative = patterns (a), (c)), and (ii) by the duration of the shift (short-term shift in the range of hours = patterns (a), (b) or long-term shifts in the range of days = patterns (c), (d)). However, pattern (e) could not be classified into any of the above categories.

Four out of 12 subjects were assigned to pattern (a) which showed an abrupt decrease in breath CH_4_ within the first few hours following vaccination. Breath CH_4_ values dropped -20 to -200 ppbv below the level of background air indicating temporary CH_4_ consumption. Following the minimum, between 12 to 24 hours after vaccine administration, CH_4_ values increased by a factor of 1 to 5 with respect to baseline values. After 1 to 4 days the base level was reached again in most of the investigated subjects. However, larger variabilities could still be observed for subjects 3 and 9 in case of the second vaccination between days 2 to 7.

Pattern (b) was found in two subjects and is characterized by a substantial positive CH_4_ peak occurring 1 to 5 hours after vaccination. For the two cases CH_4_ formation increased by a factor of ∼10 if compared with base values. In comparison to the first vaccination the CH_4_ maxima are lower for the second and third vaccination.

Two subjects assigned to pattern (c) showed decreased CH_4_ values for a period of 3 to 19 days following vaccination. Methane production was reduced by a factor of 3 to 10 compared with base values and depending on the number of vaccinations. In case of subject 5, CH_4_ values showed a sudden within the first few hours following vaccination. The largest drop can be observed in connection with the first vaccination with a stepwise decreasing tendency for subsequent vaccinations. In comparison to subject 5, subject 10 shows a prolonged decrease of CH_4_ with 100 ppbv below base level after the first vaccination whereas the second vaccination did not significantly affect CH_4_ values.

In contrast, both subjects associated with pattern (d) showed a steady increase in breath CH_4_ values after vaccine administration reaching maximum CH_4_ values of 25000 and 7000 ppbv, respectively after 1 to 4 days. Observed CH_4_ maxima following the first vaccination showed an increase by a factor of 4 to 7 compared to base levels. In case of subject 6 the second vaccination showed a similar pattern with a CH_4_ maximum of 5000 ppbv after 3 days but lower CH_4_ values when compared with first vaccination (20000 ppbv). The third vaccination showed the earliest (maximum after one day) and highest CH_4_ peak of 25000 ppbv. Interestingly, subject 2 showed reduced variations from baseline following the second and third vaccination in comparison to the immense peak following the first vaccination with a maximum after four days. The two remaining subjects 4 and 12 were categorized into (e) representing no consistent variation pattern following the three vaccinations. After the first vaccination subject 12 showed increased CH_4_ values of 35000 ppbv for a period of two days. In contrast, after the second and third vaccination CH_4_ values dropped below 10000 ppbv with a similar pattern as observed for subjects in category (c). However, CH_4_ levels increased again in the following days. Subject 4, CH_4_ values showed the most irregular pattern including extreme positive and negative CH_4_ dynamics. The measured breath CH_4_ values increased and decreased to values of 1000 ppbv and –1000 ppbv and thus by a factor of ∼100 if compared to the base range of ±10 ppbv. Following the first vaccination breath CH_4_ values even dropped to -1000 ppbv below background. In contrast, the second vaccination resulted in daily fluctuations of breath CH_4_ with decreasing maxima. However, the third vaccination showed only minor variations.

Figure 2 shows endogenous breath CH_4_ values during a COVID-19 infection of one of the authors (FK, subject 6) occurring six months after the third vaccination. Symptoms during infection were considered moderate and best described as having a “cold”. Breath CH_4_ values increased drastically after one day of being tested COVID-19 positive with a peak value of 32000 ppbv and thus by a factor of 12 higher when compared with the average CH_4_ base value of 2700 ppbv. Following the peak, breath CH_4_ production steadily decreased for a period of 3-4 days before reaching CH_4_ baseline values again. When tested COVID-19 negative, 7 days after the first positive test, CH_4_ values reflected the baseline value. It is of high interest that 28 hours prior to the first positive COVID-19 result, the CH_4_ value of 10000 ppbv was already significantly elevated compared to breath CH_4_ base values.

**Fig. 2:**
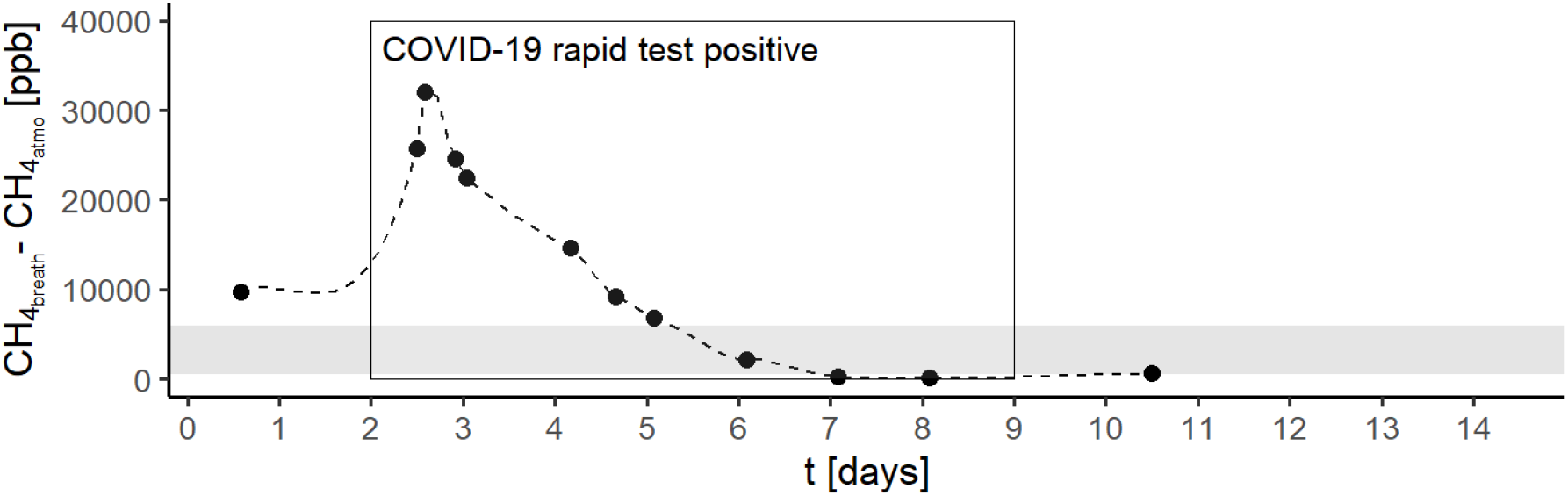
Endogenous breath CH_4 values_ (CH_4 endo_ = CH_4 breath_ – CH4 background) during COVID-19 infection. The grey shaded rectangle represents the 95% confidence interval of base CH_4 endo_ values. The black lined rectangle shows the period in which the subject was tested Covid-19 positive.

Endogenous breath CH_4_ production were reported to be closely linked to physical condition (Polag & Keppler, 2018). Figure 3 shows a comparison between physical condition and breath CH_4_ production for one of the authors (subject 6) over a time period of 74 weeks. To quantify the actual health status, the subject evaluated subjectively the physical condition on a numerical basis (scale 1 to 6) prior to breath CH_4_ measurements. Whilst 1 represents very good health conditions 6 indicates extremely bad conditions. However, 1 and 6 was not reached during the whole study period and values ranged from 1.5 to 5 with an average value of 2.5. The subjective well-being of the subject seemed to be clearly correlated to CH_4_ production with a correlation coefficient (r) of 0.47. Thus, allergic complaints (A), complaints by COVID-19 vaccination (V), a COVID-19 infection (C19) and a cold (C) occurring during the measurement period and associated with worse physical conditions (higher numbers) are accompanied by higher breath CH_4_ production (grey columns in Fig. 3).

**Fig. 3:**
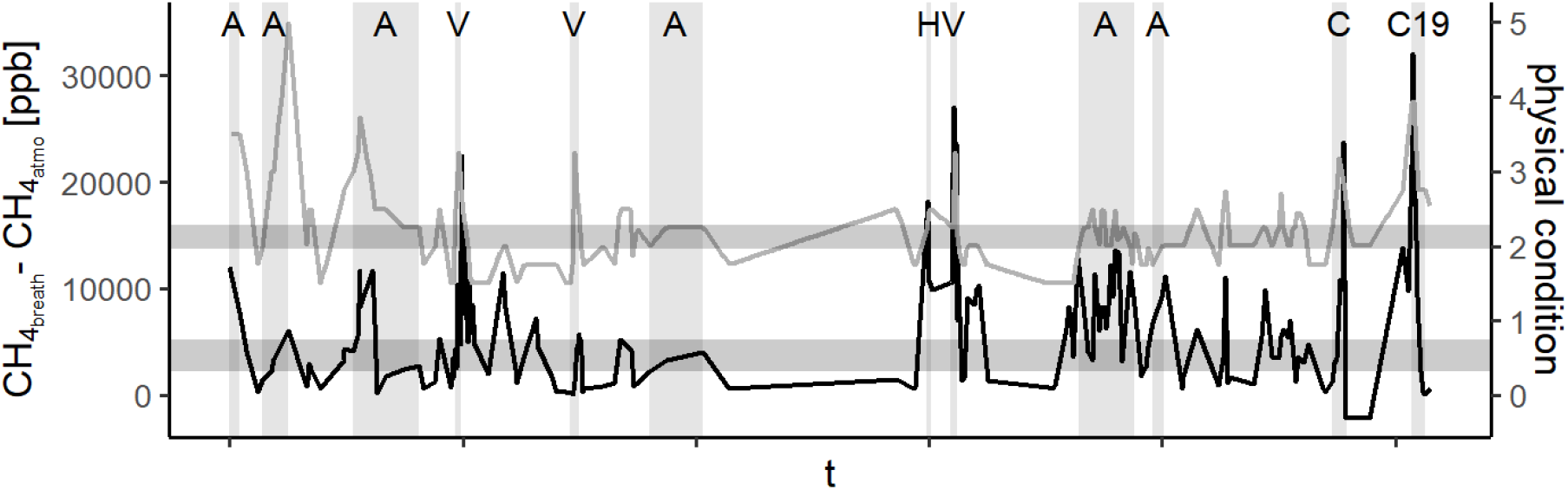
Time series of endogenous CH_4 values_ (black line) and physical conditions (grey line) of subject 6 monitored for a period of 74 weeks (r = 0.47). Horizontal grey shaded rectangles represent the 95% confidence intervals of base CH_4 endo_ values and the numerical physical condition, respectively. Vertical grey shaded columns show periods associated with worse physical conditions (A=allergic complaints, V=complaints caused by COVID-19-vaccination, H= headache, C= cold, C19=COVID-19 infection).

## 4 Discussion

We are aware that the number of studied subjects is too low to allow for broader and general conclusions. However, the limited number of subjects clearly showed an obvious effect on breath CH_4_ dynamics following COVID-19-vaccination and these patterns will be discussed as a proof of concept in closer detail in the following sections. All of the subjects showed a significant variation or shift from their individual baseline breath CH_4_ levels a few hours to days after vaccination. While intraday fluctuations of breath CH_4_ are usually in a range of 10% (Polag & Keppler, 2018), the observed breath CH_4_ values in the post-vaccination period temporally increased or decreased by a factor of 2 to 100. Most surprisingly, CH_4_ dynamics following COVID-19 vaccination seems to be characterized by an individual CH_4_ pattern which differs in (i) shift direction (positive = enhanced CH_4_ production; negative = CH_4_ consumption or reduced production) and (ii) duration of CH_4_ shift (short-term versus long-term shift). In most of the subjects the dynamic variation was more pronounced in case of the first vaccination in comparison to the second and third vaccination. The declining CH_4_ dynamic with each vaccination is most probably linked to the accumulation of immune cells and antibodies most stimulated after the first vaccination. A study by (Oberhardt, 2021) found that the cellular immune response mobilizes functional CD8^+^ T cells shortly after the first COVID-19 vaccination with an mRNA-vaccine. A relevant increase in cells was observed after six to eight days following vaccination. This period corresponds to the duration of CH_4_ shifts from baseline in most of the subjects indicating a relationship between post-vaccination CH_4_ shifts and cellular immune responses in case of COVID-19 vaccination.

Interestingly, individual post-vaccination patterns were also observed after the administration of other vaccines and during a COVID-19 infection (Fig. 2). Subject 5 showed a similar pattern (i.e., long-term decrease of CH_4_; pattern (c) in Fig. 1) after a first-time tick-borne encephalitis vaccination a few months prior to COVID-vaccination (data not shown). Analogous, subject 6 showed a long-term increase in CH_4_ subsequent to a first-time typhus vaccination (data not shown).

Potential intrinsic host factors correlating with the observed post-vaccination CH_4_ pattern (Fig. 1) are age and CH_4_ emission status. Short-term patterns where post-vaccination CH_4_ values dropped back to baseline after a period of 1 to 2 days were usually observed in connection with a lower average age and (mean: 27 years, n=6) /or a low CH_4_ emission status (5 low, 1 high). In contrast, long-term patterns with a period of 14 days are more frequently observed in connection with a higher average age (mean: 47 years, n=4) and/or median to high CH_4_ emission status (3 medium/high, 1 low). Both factors are possibly interrelated as age is a dominant factor which affects CH_4_ emission status (Polag, et al., 2014). Generally, age is known to have a major influence on immune responses to vaccination (Zimmermann & Curtis, 2019). Aging is accompanied by an increasing imbalance between pro-inflammatory and anti-inflammatory cytokines (Rea, et al., 2018). Inflammaging is associated with an increase in pro-inflammatory cytokines causing a decline in CD8^+^ T cells, and thus, resulting in a reduction of immune responses with respect to inflammation processes. Post-vaccination fluctuations in breath CH_4_ might be indicating the maintenance of redox homeostasis in cells. Short-term fluctuations directly following vaccination might also occur in subjects with long-term dynamics but are probably overlaid by the higher baseline level of CH_4_.

The post-vaccination dynamics in breath CH_4_ suggest that CH_4_ might be an indicator of individual immune responses following the induced immune perturbation by vaccination. Interestingly, not only an induced immune perturbation leads to a shift in CH_4_ production but also ‘natural’ immune perturbation seems to affect CH_4_ dynamics. Subject 6 (FK) was tested COVID-19 positive around half a year after his third vaccination and sampled breath air before, whilst and after infection being tested positive (Fig. 2). Methane production during COVID-19 infection showed a similar pattern to CH_4_ dynamics following the first, second and third COVID-19 vaccination (pattern (d) in Fig. 1). However, in comparison to the vaccinations, the COVID-19 infection showed the largest deviations from baseline CH_4_ values. Moreover, the CH_4_ production was already enhanced before COVID-19 rapid test was positive. Thus, if the breath CH_4_ base line of an individual subject is known, sudden significant shifts during a frequent monitoring of breath CH_4_ values might indicate a deviation from the state of health. This is also confirmed by the correlation between periods of physical discomfort caused by inflammation (immune perturbation, infections and allergic reactions), and enhanced breath CH_4_ values (Fig. 3). A previous breath CH_4_ monitoring study by (Polag & Keppler, 2018) also showed that a sudden temporary increase in endogenous CH_4_ was often linked to immune reactions and inflammatory processes.

The underlying processes behind temporary increased formation of endogenous CH_4_ and CH_4_ consumption following immune reactions are not yet completely understood. However, increased CH_4_ production might be explained by a just discovered non-enzymatic mechanism of endogenous CH_4_ production in cells that has recently been proposed by (Ernst, et al., 2022). Based on laboratory experiments with living cells from all domains of life including several human cell lines the authors revealed a mechanism which is based on reactive oxygen species (ROS), free iron and methyl groups bound to nitrogen and sulphur as for example occurring in compounds such as methionine, dimethyl sulfoxide and trimethylamine. Thus, an elevated level of ROS by enhanced oxidative stress and redox regulation during immune reactions and inflammatory processes leads to increased formation of CH_4_. As vaccination represents an induced perturbation of the immune system accompanied by enhanced oxidative stress, the results from this study give evidence that the described mechanism might also occur in human cell physiology.

However, it cannot be completely ruled out that the observed temporary increase in breath CH_4_ was produced by microbes. Alternatively, the methanogenic population in the gut might somehow be influenced by the immune response due to vaccination. Recent studies found that the human microbiome has a key role in the modulation of the human immune system (Ifeanyi, 2018). Generally, the impact of microbiota as a factor that influences the immune response to vaccination has been little considered so far, but is of immense importance (Zimmermann & Curtis, 2019). (Hagan, et al., 2019) observed that antibiotics treatment associated with a loss in gut microbial diversity altered vaccine immunity in subjects with low pre-existing immunity. Inversely, the vaccine itself while stimulating the immune system might have an effect on quality and quantity of the gut microbiome. Thus, increased breath CH_4_ formation following vaccination might derive either by stimulating methanogenic growth and activity or by inhibiting methanogenesis by alternative pathways for example by an increase in sulphate reducing bacteria or acetogenic bacteria as competitors for hydrogen (H_2_) (Christl, et al., 1992), (Sahakian, et al., 2010).

However, post-vaccination endogenous breath CH_4_ values falling below those of ambient air (inhaled air) which were observed for low CH_4_ breath emitters can only be explained by CH_4_ degradation processes. To our knowledge, CH_4_ consumption in the human body has not been reported before. Thus, hypotheses on potential chemical or microbial degradation pathways are missing so far. Methanotrophic bacteria or archaea which are capable of utilizing CH_4_ as sole carbon and energy source can be found in a wide range of environments such as groundwater, rice fields or the gut of termites (Conrad, et al., 2009). However, the existence of methanotrophs in the human gut has not been described so far but recent estimates indicate that 40-50% of human gut species not yet genomically identified (Nayfach, et al., 2019). Thus, in principle it is possible that a strain of bacteria or archaea known to oxidize CH_4_ also resides in the human gut and leads to the observed temporary loss in endogenous breath CH_4_ triggered by immune reactions. Chemical degradation of CH_4_ analogous to CH_4_ removal by hydroxyl radicals in the atmosphere might also explain CH_4_ consumption as OH-radicals are permanently formed as byproducts during metabolism. Under natural conditions CH_4_ interacts with OH-radicals and forms a reaction chain including a series of organic compounds such as methyl (CH_3_) and formaldehyde (CH_2_O) (Lu & Khalil, 1993), (Dzyuba, et al., 2012).

However, whether the observed CH_4_ dynamics following immune reactions are of enzymatic or chemical nature, our study highlights the great potential of CH_4_ as a biomarker as also recently discussed by (Wang, 2014), (Boros & Keppler, 2019) and (Keppler, et al., 2022). First, breath CH_4_ seems to be an indicator of increased hypoxic events and redox regulation during immune reactions/inflammatory processes. This is confirmed by breath CH_4_ deviations from baseline observed in case of infections/immune reactions accompanied by physical discomfort. Second, CH_4_ might be a marker of individual immune responses or immune states. This is confirmed by individual CH_4_ variation patterns following COVID-19 vaccination.

In future medicine it is of great interest to improve vaccine immunogenicity and efficacy (Tsang, et al., 2020). Therefore, it is necessary to define individual immune states (Shen-Orr & Furman, 2013). As a vaccine represents an induced perturbation of the immune system, it is an adequate method to study human immune response. As many factors influence the immune state including intrinsic factors (age, sex, genetics) and vaccine factors (vaccine type etc.) as dominant factors beyond many others (nutritional, environmental, etc.) it would be of great benefit to have an in vivo parameter which classifies the efficacy of a specific vaccine. Therefore, in future studies it would be necessary to record additional immune parameters such as antibody level to calibrate breath CH_4_ values in terms of quality and quantity of immune responses.

## 5 Conclusions

This study should been considered as a proof of principle that breath CH_4_ dynamics in humans are affected by cellular CH_4_ formation and degradation processes. Following vaccination as an induced perturbation of the immune system we reported the first indications that significant deviations (both positive and negative) from average breath CH_4_ values are potentially related to immune reactions and might also originate from redox homeostasis in cells. The observed individual breath CH_4_ dynamics further indicate that CH_4_ has a potential bioactive role in immunology as already suggested by (Boros & Keppler, 2019). A change in breath CH_4_ level from baseline values might be used to detect changes in individual ROS levels and oxidative stress and could possibly classify immune responses. Thus, in in the field of system biology and precision medicine, breath CH_4_ might be used as a diagnostic tool. For more details regarding the role of oxidative stress in cellular CH_4_ formation and its potential significance for clinical and health sciences we would like to refer the readers to the article by (Keppler, et al., 2022). Follow up studies should focus on deciphering the potential physiological role of CH_4_ in humans including monitoring CH_4_ as an oxidative stress biomarker and indicator of individual immune states. The advantages of CH_4_ compared to conventional breath biomarkers is its easy detection by analytical methods such as GC-FID and laser spectroscopy (Keppler, et al., 2016). Thus, no elaborate or costly sample analysis is necessary (Risby & Sehnert, 1999). However, further investigations are necessary to provide clear evidence of non-microbial CH_4_ production in humans and the underlying processes of its formation and consumption. This will be a considerable challenge since CH_4_ production by methanogens is the dominant process in case of high emitters and will make it difficult to distinguish the microbial pathway of CH_4_ production from the non-microbial pathway.

## Acknowledgment

We would like to thank the Biogeochemistry group at the Institute of Earth Sciences/Heidelberg University for providing breath methane samples. Financial support by the German Science Foundation (DFG KE 884/8-2).

## References

Aronson, J. K. & Ferner, R. E., 2017. Biomarkers - A general review. Current Protocols in Pharmacology, Band 76.

Bang, C. et al., 2014. The intestinal archaea Methanospaera stadtmanae and Methanobrevibacter smithii activate human dendritic cells. PLoS One, Band 9, pp. 1–9.

Bolin, T. D., Myo-Khin, S.-A., Genge, J. R. & Duncombe, V. M., 1996. Correlation of hydrogen and methane production to rice carbohydrate malabsorption in Burmese (Myanmar) children. Journal of Pediatric Gastroenterology and Nutrition, Band 22, pp. 144–147.

Bond, J. H., Engel, R. R. & Levitt, M. D., 1971. Factors influencing pulmonary methane excretion in man. The Journal of Experimental Medicine, Band 133, pp. 572–588.

Boros, M. & Keppler, F., 2018. Gasotransmitters. In: R. Wang, Hrsg. s.l.:The Royal Society of Chemistry, pp. 192–234.

Boros, M. & Keppler, F., 2019. Methane production and bioactivity-A link to oxido-reductive stress. Frontiers in Physiology.

Boros, M., Tuboly, E., Meszaros, A. & Amann, A., 2015. The role of methane in mammalian physiology-is it a gasotransmitter?. Journal of Breath Research, Band 9.

Christl, S. U., Gibson, G. R. & Cummings, J. H., 1992. Role of dietary sulphate in the regulation of methanogenesis in the human large intestine. Gut, Band 33, p. 1234–1238.

Conrad, R., Klose, M. & Noll, M., 2009. Functional and structural response of the methanogenic microbial community in rice field soil to temperature change. Environmental Microbiology, Band 11, p. 1844–1853.

Conway de Macario, E. & Macario, A. J. L., 2009. Methanogenic archaea in health and disease: A novel paradigm of microbial pathogenesis. International Journal of Medical Microbiology, Band 299, pp. 99–108.

Das, S. & Pal, M., 2020. Review—Non-invasive monitoring of human health by exhaled breath analysis: A comprehensive review. Journal of the Electrochemical Society, Band 167.

De Lacy Costello, B. P. J., Ledochowski, M. & Ratcliffe, N. M., 2013. The importance of methane breath testing: a review. Journal of Breath Research, Band 7.

Di Stefano, M. & Corazza, G. R., 2009. Role of hydrogen and methane breath testing in gastrointestinal diseases. Digestive and Liver Disease Supplements, Band 3, pp. 40–43.

Dryahina, K., Smith, D. & Spanel, P., 2010. Quantification of methane in humid air and exhaled breath using selected ion flow tube mass spectrometry. Rapid Communications in Mass Spectrometry, Band 24, pp. 1296–1304.

Dzyuba, A. V., Eliseev, A. V. & Mokhov, I. I., 2012. Estimates of changes in the rate of methane sink from the atmosphere under climate warming. Atmospheric and Oceanic Physics, Band 48, p. 332–342.

Ernst, L. et al., 2022. Methane formation driven by reactive oxygen species across all living organisms. Nature.

Flourie, B. et al., 1991. Site and substrates for methane production in human colon. Gastrointestinal and Liver Physiology, Band 260, pp. 752–757.

Furnari, M. et al., 2012. Reassessment of the role of methane production between irritable bowel syndrome and functional constipation. Journal of Gastrointestinal and Liver Diseases, Band 21, pp. 157–163.

Gaude, E. et al., 2019. Targeted breath analysis: exogenous volatile organic compounds (EVOC) as metabolic pathway-specific probes. Journal of Breath Research, Band 13.

Ghavami, S. B. et al., 2018. Alterations of the human gut Methanobrevibacter smithii as a biomarker for inflammatory bowel diseases. Microbial Pathogenesis, Band 117, pp. 285–289.

Ghosh, C. et al., 2021. Breath-based diagnosis of infectious diseases: A review of the current landscape. Clinics in Laboratory Medicine, Band 41, pp. 185–202.

Ghyczy, M. et al., 2008. Hypoxia-induced generation of methane in mitochondria and eukaryotic cells: an alternative approach to methanogenesis. Cell physiology and Biochemistry, Band 21, pp. 251–258.

Hagan, T. et al., 2019. Antibiotics-driven gut microbiome perturbation alters immunity to vaccines in humans. Cell, Band 178, p. 1313–1328.

Hayney, M. S. & Buck, J. M., 2002. Effect of age and degree of immune activation on cytochrome P450 3A4 activity after influenza immunization. Pharmacotherapy, Band 22.

Hong-Geller, E. & Adikari, S., 2018. Biosensing technologies for the detection of pathogens – a prospective way for rapid analysis. In: R. Rinken & K. Kivirand, Hrsg. s.l.:IntechOpen, pp. 21–36.

Hopkins, M. J., Sharp, R. & Macfarlane, G. T., 2002. Variation in human intestinal microbiota with age. Digestive Liver Disease, Band 34, pp. 512–518.

Hwang, L. et al., 2010. Evaluating breath methane as a diagnostic test for constipation-predominant IBS. Digestive Diseases and Sciences, Band 55, pp. 398–403.

Ifeanyi, O. E., 2018. A review on free radicals and antioxidants. International Journal of Current Research in Medical Sciences, Band 4, p. 123–133.

Keppler, F. et al., 2009. Methane formation in aerobic environments. Environmental Chemistry, Band 6, pp. 459–465.

Keppler, F., Hamilton, J. T. G., Brass, M. & Röckmann, T., 2006. Methane emissions from terrestrial plants under aerobic conditions. Nature, Band 439.

Keppler, F. et al., 2016. Stable isotope and high precision concentration measurements confirm that all humans produce and exhale methane. Journal of Breath Research, Band 10.

Kinoyama, M. et al., 2006. Diurnal variation in the concentration of methane in the breath of methane producers. Microbial Ecology in Health and Disease, Band 18, pp. 47–54.

Kunkel, D. et al., 2011. Methane on breath testing is associated with constipation: A systematic review and meta-analysis. Digestive Diseases and Sciences, Band 56, pp. 1612–1618.

Lenhart, K. et al., 2012. Evidence for methane production by saprotrophic fungi. Nature Communications, Band 3, p. 1046.

Lenhart, K. et al., 2016. Evidence for methane production by the marine algae \textitEmiliana huxleyi. Biogeosciences, Band 13, pp. 3163–3174.

Lenhart, K. et al., 2015. Nitrous oxide and methane emissions from cryptogamic covers. Global Change Biology, Band 21, pp. 3889–3900.

Lu, Y. & Khalil, M. A., 1993. Methane and carbona monoxide in OH chemistry: The effects of feedbacksand reservoirs generated by the reactive products. Chemosphere, Band 26, p. 641–655.

Manolis, A., 1983. The diagnostic potential of breath analysis. Clinical Chemistry, Band 29, pp. 5–15.

McKay, L. F., Eastwood, M. A. & Brydon, W. G., 1985. Methane excretion in man - a study of breath, flatus, and faeces. Gut, Band 26, pp. 69–74.

Mihajlovski, A. et al., 2010. Molecular evaluation of the human gut methanogenic archaeal microbiota reveals an age-associated increase of the diversity. Environmental Microbiology Reports, Band 2, pp. 272–280.

Montes, R. G., Saavedrea, J. M. & Perman, J. A., 1993. Relationship between methane production and breath hydrogen excretion in lactose-malabsorbing individuals. Digestive Diseases and Sciences, Band 38, pp. 445–448.

Mürtz, M. & Hering, P., 2008. Mid Infrared Coherent Sources and Applications. In: M. Ebrahim-Zadeh & I. T. Sokorina, Hrsg. s.l.:Springer, pp. 535–555.

Nakhleh, M. K. et al., 2017. Diagnosis and Classification of 17 Diseases from 1404 Subjects via Pattern Analysis of Exhaled Molecules. {ACS} Nano, jan, 11(1), pp. 112–125.

Nayfach, S. et al., 2019. New insights from uncultivated genomes of the global human gut microbiome. Nature, Band 568, p. 505–510.

Oberhardt, V. e. a., 2021. Rapid and stable mobilization of CD8+ T cells by SARS-CoV-2 mRNA vaccine. Nature, Band 597.

Peled, Y., Weinberg, D., Hallak, A. & Gilat, T., 1987. Factors affecting methane production in humans. Digestive Diseases and Sciences, Band 32, pp. 267–271.

Phillips, M. et al., 2010. Effect of influenza vaccination on oxidative stress products in breath. Journal of Breath Research, Band 4.

Pitt, P. et al., 1980. Studies on breath methane: The effect of ethnic origins and lactulose. Gut, Band 21, pp. 951–959.

Polag, D. & Keppler, F., 2018. Long-term monitoring of breath methane. Science of The Total Environment, may, Band 624, pp. 69–77.

Polag, D., Leiß, O. & Keppler, F., 2014. Age dependent breath methane in the German population. Science of the Total Environment, Band 481, pp. 582–587.

Pulendran, B. & Davis, M., 2020. The science and medicine of human immunology. Science, Band 369.

Rea, I. M. et al., 2018. Age and age-related diseases: Role of inflammation triggers and cytokines. Frontiers in Immunology, Band 9.

Risby, T. H. & Sehnert, S. S., 1999. Clinical application of breath bbiomarker of oxidative stress status. Free Radical Biology & Medicine, Band 27, p. 1182–1192.

Roccarina, D. et al., 2010. The role of methane in intestinal diseases. Clinical and Systematic Reviews, Band 105, pp. 1250–1256.

Sahakian, A., Jee, S.-R. & Pimentel, M., 2010. Methane and the gastrointestinal tract. Digestive Diseases and Sciences, Band 55, p. 2135–2143.

Santos-Mello, C. et al., 2012. Methane production and small intestinal bacterial overgrowth in children living in a slum. World Journal of Gastroenterology, Band 18, pp. 5932–5939.

Shen-Orr, S. S. & Furman, D., 2013. Variability in the immune system: of vaccine responses and immune states. Current Opinion in Immunology, Band 25, p. 542–547.

Szabó, A. et al., 2015. Exhaled methane concentration profiles during excercise on an ergometer. Journal of Breath Research, Band 9.

Triantafyllou, K., Chang, C. & Pimentel, M., 2014. Methanogens, methane and gastrointestinal motility. Journal of Neurogastroenterology and Motility, Band 20, pp. 31–40.

Tsang, J. S. et al., 2020. Improving vaccine-induced immunity: Can baseline predict outcome?. Trends in Immunology, Band 41, p. 457–465.

Tuboly, E. et al., 2017. Excessive alcohol consumption induces methane production in humans and rats. Scientific Reports, Band 7, p. 7329.

Tuboly, E. et al., 2013. Determination of endogenous methane formation by photoacoustic spectroscopy. Journal of Breath Research, Band 7.

Wang, R., 2014. Gasotransmitters: Growing pains and joys. Trends in Biochemical Sciences, Band 5, p. 227–232.

Zimmermann, P. & Curtis, N., 2019. Factors that influence the immune response to vaccination. Clinical Microbiology Reviews, Band 32.

